# Expansion microscopy resolves the 3D thylakoid structure

**DOI:** 10.1101/2023.05.17.541202

**Authors:** Peter R. Bos, Jarne Berentsen, Emilie Wientjes

## Abstract

The light-harvesting reactions of photosynthesis take place on the thylakoid membrane inside chloroplasts. The thylakoid membrane is folded into appressed membranes, the grana, and non-appressed membranes that interconnect the grana, the stroma lamellae. This folding is essential for the correct functioning of photosynthesis. Electron microscopy and atomic force microscopy are commonly used to study the thylakoid membrane, but these techniques have limitations in visualizing a complete chloroplast and its organization. To overcome this limitation, we applied expansion microscopy (ExM) on isolated chloroplasts. ExM is a technique that involves physically expanding a sample in a swellable hydrogel to enhance the spatial resolution of fluorescence microscopy. Using all-protein staining, we have visualized the 3D structure of spinach thylakoids with a high level of detail. We were able to resolve stroma lamellae that were 60 nm apart and observe their helical wrapping around the grana. Furthermore, we accurately measured the dimensions of grana from top-views of chloroplasts, which allow for precise determination of the grana diameter. Ultimately, we constructed a 3D model of a complete chloroplast, which provides a foundation for structure-based modeling of photosynthetic adaptations. Our results demonstrate that ExM is a fast and reliable technique for studying thylakoid organization with a high level of detail.

## Introduction

Photosynthesis powers virtually all life on Earth. The initial steps of photosynthesis, light-harvesting and electron transfer, take place in the thylakoid membrane, a single continuous membrane located in the chloroplast (Blankenship, 2021). The thylakoid membrane is intricately folded into non-appressed membranes, called stroma lamellae, and appressed membrane stacks, called grana. The grana stacks are approximately cylindrical with a diameter of 280-600 nm, but this diameter is variable in response to different light conditions (Mehta et al., 1999; Kaftan et al., 2002; Shimoni et al., 2005; Fristedt et al., 2009; Anderson et al., 2012; Armbruster et al., 2013; Pribil et al., 2014; Schumann et al., 2017; Wood et al., 2018; Bussi et al., 2019; Sattari Vayghan et al., 2022). The stroma lamellae are wrapped around the grana in a right-handed helix and are connected to the grana at slit-like apertures (Bussi et al., 2019). The thylakoid membrane has a protein concentration of 70%, but remains highly dynamic (Kirchhoff, 2014). Moreover, diffusion within the membrane and between the grana and stroma is fast, on the sub-second timescale for plastocyanin and still on the minute timescale for protein complexes (Kirchhoff, 2014; Höhner et al., 2020). How the thylakoid, with its complex 3D architecture can be so dynamic and what factors facilitate it, is not well understood (Johnson and Wientjes, 2020).

The folding and organization of the thylakoid membrane have been extensively studied using various techniques (Pribil et al., 2014; Blankenship, 2021). Electron microscopy (EM) is the most commonly used method, with transmission EM (TEM) providing sufficient resolution to image single membrane bilayers within each granum stack. TEM images have revealed the helical wrapping of stroma around the grana (Paolillo Jr, 1970; Mustárdy and Garab, 2003; Mustardy et al., 2008). A further increase in resolution has been achieved using scanning EM and focused ion beam scanning EM, which showed a left-handed helix in the stroma lamellae as well (Bussi et al., 2019). Lastly, Cryo-EM has been used to study the thylakoid membrane of *Chlamydomonas reinhardtii* with nanometer resolution (Engel et al., 2015; Wietrzynski et al., 2020). However, constructing a 3D model of the entire thylakoid membrane using EM is challenging due to the need for thin sample slices (max 200 nm) and the time-consuming and expensive sample preparation and imaging (Wassie et al., 2019). Additionally, localizing specific proteins in the thylakoid membrane is difficult, since many of the proteins hardly protrude from the membrane (Johnson et al., 2014; Wietrzynski et al., 2020). Specific protein localization is possible with atomic force microscopy, but this technique can image only a single layer of membrane (Liu and Scheuring, 2013; Wood et al., 2018; Onoa et al., 2020). Fluorescence microscopy can resolve the position of the grana but lacks the resolution to study the membrane (Mehta et al., 1999; Wildman et al., 2005). Although Structure Illuminated Microscopy (SIM) has improved the resolution of fluorescence microscopy, it still lacks the desired molecular detail (Iwai et al., 2018; Wood et al., 2019; Flannery et al., 2021). Other single molecule or super resolution microscopy techniques have not yet been applied on the thylakoid membrane, mainly due to difficulties with the massive autofluorescence of chlorophyll in such a complex system (Johnson and Wientjes, 2020). Thus, while imaging whole chloroplasts with fluorescence microscopy is fast and easy, the resolution of these techniques is not sufficient to resolve the fine structures of the thylakoid membrane. Hence, chloroplasts are too large to be easily visualized with EM but too small to be accurately imaged with fluorescence microscopy.

The gap in resolution between EM and fluorescence microscopy has been bridged with the introduction of expansion microscopy (ExM). In ExM, a sample is physically expanded isotropically in a swellable hydrogel, resulting in a larger distance between fluorophores and proteins (Chen et al., 2015; Gambarotto et al., 2019; Damstra et al., 2022). By doing so, the effective resolution of the sample is improved. Moreover, the sample is de-crowded, which increases diffusion and epitope recognition by antibodies (Chen et al., 2015). Several methods have been developed to stain lipids, all proteins, and/or only specific proteins (Damstra et al., 2022).

In this work, we developed a method for ExM on de-enveloped chloroplasts. We combined ultrastructure-ExM (U-ExM), an optimized version of the original protein-retention ExM protocol, with an all-protein staining (Pan-ExM) (Gambarotto et al., 2019; M’Saad and Bewersdorf, 2020) and achieved a 4.8-6.7 times expansion of chloroplasts. We measured or quantified the dimensions of the grana and confirmed the right-handed helical stroma around the grana. Together this shows that chloroplasts can be imaged easily, fast, and accurately and presents the potential of using ExM in future studies to reveal the dynamics of the thylakoid membrane.

## Material and Methods

### Isolation of de-enveloped chloroplasts

*Spinacia oleracea* (Spinach) was purchased from the local grocer. Chloroplasts were isolated in the dark according to an adapted protocol from Caffari et al. (Caffarri et al., 2009). In short, leaves were chilled in ice water. They were then quickly homogenized in a blender in ice cold buffer 1 (B1: 400 mM sorbitol, 5 mM EDTA, 10 mM NaHCO_3_, 5 mM MgCl_2_ and 20 mM tricine). The resulting suspension was filtered through a cheesecloth and the (de-enveloped) chloroplasts were pelleted by centrifugation (1500 ×g, 5 min, 4 ᵒC). The pellet was carefully resuspended in buffer 1. Centrifugation and resuspension were repeated twice. Chloroplasts were kept at 4 ᵒC in the dark until further use.

### Chloroplast fixation

The protocol for (de-enveloped) chloroplast fixation was based on the fixation of mitochondria as described by Gambarotto and co-workers (Gambarotto et al., 2019). For the fixation, paraformaldehyde (PFA) was warmed to 37 ᵒC for 30 minutes to increase its reactivity. Chloroplasts were fixed in a 3% PFA, 0.1% glutaraldehyde (GA) solution in B1 for 30 minutes at room temperature. They were then washed twice in B1 and permeabilized in 0.1% Triton X-100 in B1 for 3 minutes on ice. After three wash steps, chloroplasts were anchored in a solution of 1.0% acrylamide (AA) and 0.7% PFA overnight at room temperature in the dark. The anchor of AA is essential for covalent bonding of the proteins to the sodium acrylate (SA) - AA gel. Chloroplasts were washed 4 times to remove traces of PFA and AA and stored at 4 ᵒC until further use. Chloroplasts retained their structure for at least two weeks when kept cool and in the dark.

### Gel composition

SA was synthesized according to the protocol from Damstra et al. (Damstra et al., 2022) or purchased from Sigma-Aldrich. The gel composition was based on the Ultrastructure-ExM protocol from Gambarotto and co-workers (Gambarotto et al., 2019).

Several gel compositions with different expansion factors have been used to test expansion factor of the gel and expansion factor of the chloroplasts (table 1). All gel compositions contained 1.1× phosphate buffered saline (PBS). Gel composition B has been used for most images in this article.

**Table 1.** Composition of four gels used in this study. All percentages are w/v%. For expansion factor, see SI figure 2

|  | Sodium acrylate | Acrylamide | N,N'-methylenebisacrylamide |
| --- | --- | --- | --- |
| Gel A | 20.9% | 10.0% | 0.10% |
| Gel B | 30.1% | 10.8% | 0.04% |
| Gel C | 25.4% | 5.6% | 0.01% |
| Gel D | 34.3% | 5.3% | 0.01% |

### Chloroplast expansion

Fixed and anchored chloroplasts were mixed with the gel solution in a 1:10 ratio and kept on ice. Gel polymerization was started by adding 0.1 w/v% N,N,N′,N′-tetramethyl ethylenediamine (TEMED) and 0.1 w/v% ammonium persulphate (APS) from 10 w/v% stocks, and the solution was quickly pipetted in a polymerization chamber as described by Zhang et al. (Zhang et al., 2020). Alternatively, a drop of fixed and anchored chloroplasts was spread on a coverslip and allowed to dry for 20 minutes. Again, gel polymerisation was started by adding TEMED and APS at 0.1 w/v%. The solution was quickly pipetted in a polymerization chamber and closed with the coverslip with the layer of chloroplasts in the gel solution.

The gels were allowed to polymerize for 1.5 h in a humid environment at room temperature. Afterwards, they were removed from the chamber, cut into an asymmetrical shape, photographed and expanded in a petri dish in ultrapure water. After 3 to 5 washing steps in ultrapure water, full expansion was achieved and the gel was photographed again. The expansion factor of the gel was determined dividing the dimensions of the gel after expansion by its dimensions before expansion. A small piece was cut out and put in an 8-well plate. The gel was washed twice in 0.1 M NaHCO3, pH 8.3 and stained in 20 μg/mL N-Hydroxysuccinimide (NHS) ester-ATTO488 (ATTO-TEC GmbH, Art. Nr.: AD 488) in 0.1 M NaHCO3, pH 8.3 for 1.5 h. After staining, the gel was washed several times in ultrapure water to achieve full expansion and remove any unbound staining.

### Imaging

Unexpanded chloroplasts - A few microliter of fixed chloroplasts were placed on a microscope slide and covered with a coverslip. The chloroplasts were imaged with a confocal TCS SP8 system from Leica Microsystems equipped with an HC PL APO CS2 63×/1.20 NA water immersion objective and a white light laser. Excitation wavelength was set to 620 nm and detection wavelength to capture chlorophyll fluorescence (670-730 nm). Z-stacks were recorded to image the complete chloroplasts.

ExM imaging - The gels were imaged with one of several microscopes. We used a confocal TCS SP8 system from Leica Microsystems equipped with a HC PL APO CS2 63×/1.20 NA water immersion objective and an argon laser. Alternatively, we used a confocal TCS SP8 system from Leica Microsystems equipped with a HC PL IRAPO 40×/1.10 NA water immersion objective and two-photon excitation. Lastly, we used the ZEISS Elyra 7 with Lattice SIM² with a C-Apochromat 63×/1.2 NA water immersion objective at the ZEISS demo-center in Oberkochen. Excitation was set to 488 nm (single photon excitation) or 750 nm (two-photon excitation) and emission to only record ATTO488 signal (505-540 nm). Z-stacks were recorded to image complete chloroplasts. A novel image reconstruction algorithm from ZEISS was applied on the images from the Elyra 7 with Lattice SIM². All three microscopes recorded mirror images, so the recorded images were mirrored back before image analysis.

### Image analysis

Chloroplast dimensions and expansion factor - To estimate the expansion factor, we imaged chloroplasts before and after expansion. Only top-view images of chloroplasts were used to measure their size. Unexpanded chloroplasts and expanded chloroplasts were detected in each slice of the Z-stack and measured by a custom-written FIJI script. This script returned for each detected chloroplast measures like area, mean intensity and circularity as calculated by 4π*area/perimeter^2. Only objects that met set criteria for circularity and area were selected. The shape of a chloroplast was assumed to be circular and the radius was calculated from the area. Moreover, an ellipse was fitted around the object and the X and Y coordinates of the center were returned. Based on these coordinates, chloroplasts that appeared in multiple images of the z-stack were clustered by a custom-written Python script. The script to detect chloroplasts could make mistakes in a few slices of the images, for example by grouping neighboring chloroplasts. We set the minimal number of slices in which a chloroplast needed to be detected to 4, to prevent these outliers from appearing in the data. The 75^th^ percentile was taken as the size of the chloroplasts. This value was on average about 95% of the maximum value. The expansion factor of the chloroplasts was calculated by dividing the average radius of chloroplasts in a single gel by the average radius of unexpanded and fixed chloroplasts.

Grana dimensions - The grana diameter was determined with custom written scripts in FIJI, Google Colab and Python. Top-view images of chloroplasts with clear grana were selected and integrated to have a the same pixel size in pre-expansion dimensions. Grana were detected by a the Stardist 2D plugin (Schmidt et al., 2018; Weigert et al., 2020; Gómez-de-Mariscal et al., 2021). Stardist is a machine learning tool to detect convex-star shaped objects. A Stardist model was trained on an image set of chloroplast images with annotated grana. FIJI was used to measure the ROIs generated by Stardist in the original image and Python was used to cluster detected grana with similar X, Y and Z coordinates. The maximum detected size of a clustered grana value set was taken. The values were divided by the expansion factor as measured from chloroplast expansion to retrieve the pre-expansion size of the grana.

Grana height was measured by hand in FIJI in side-view images from chloroplasts. Values were divided by the expansion factor as measured from chloroplast expansion to retrieve the pre-expansion dimensions.

The number of grana was counted by hand in FIJI in 20 randomly selected images containing top views of chloroplasts.

Resolution – The minimal distance that could be distinguished with ExM was determined by making intensity profiles in images with stroma lamellae in close proximity. The full-width half maximum (FWHM) was determined and its middle was taken as the center of the peak. Distance between the peaks was calculated in pre-expansion dimensions.

All scripts and models are made available on Github (https://git.wur.nl/peter1.bos/230322-exm-script-for-chloroplast-and-grana-detection.git).

### 3D reconstruction

3D reconstruction was performed with Drishti (Limaye, 2012). First, FIJI was used to smooth a top-view image and integrate it in X, Y and Z to get a voxel size of 60 nm in all directions. Drishti Paint was then used to segment grana from the rest of the image (Hu et al., 2020). The grana segment was colored differently than the surroundings and a 3D reconstruction of a chloroplast and its grana was animated.

## Results

The thylakoid structure of chloroplasts from plants has been investigated with various imaging techniques, such as EM, AFM and confocal microscopy (Pribil et al., 2014; Wietrzynski et al., 2020; Blankenship, 2021). However, there is a resolution gap among these techniques that can be bridged using ExM. In this study, we developed a method for ExM on isolated chloroplasts. The chloroplasts were fixed and permeabilized to maintain the thylakoid structure and make the chloroplasts less resistant to expansion. Most of the chlorophyll was washed away during the permeabilization step. Proteins in the sample were linked to acrylamide anchors, which covalently link to the sodium acrylate-acrylamide gel in the gelation step (figure 1A) (Lai et al., 2016; Gambarotto et al., 2019). The primary amines of lysines and N-termini were labelled with an ATTO-488-NHS-ester staining, after which the samples were expanded 4.8 to 6.7 times and imaged. We found that the chloroplasts expanded with the gel up to a gel expansion factor of 6. However, chloroplasts expanded less than the gel when the gel expansion was higher (up to 10 times, SI figure 1 and 2). We observed a clear increase in resolution and could easily identify individual grana in the expanded chloroplasts (figure 1B). A great advantage of ExM over EM-and AFM-based methods is that a z-stack of an entire chloroplast can be recorded to resolve the complete thylakoid structure of single chloroplasts in 3D (figure 2 and SI movie 4).

**Figure 1.**
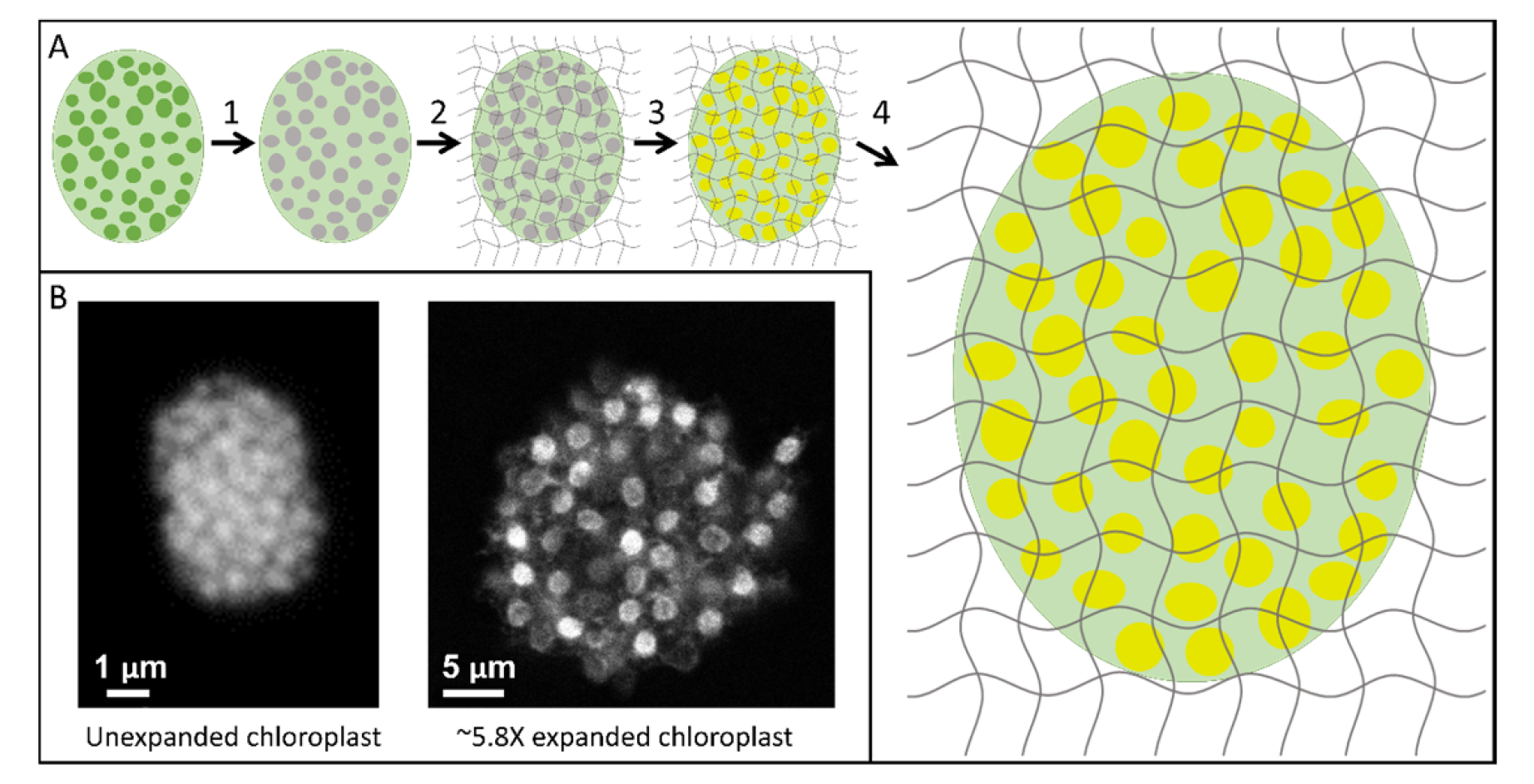
ExM on chloroplast. A) Workflow of ExM on chloroplasts. Isolated chloroplasts were fixed, permeabilised and anchored, upon which Chl fluorescence was lost (1). The chloroplasts were put in gel solution (2), stained with an ATTO-488 NHS-ester staining to stain all proteins (3) and expanded in ultrapure water (4). B) Chlorophyll fluorescence of an isolated unexpanded chloroplast (left) and ATTO-488 fluorescence of a 5.8 times expanded chloroplast (right). Both images made with confocal microscopy. Scalebars represent distance without correction for expansion.

**Figure 2.**
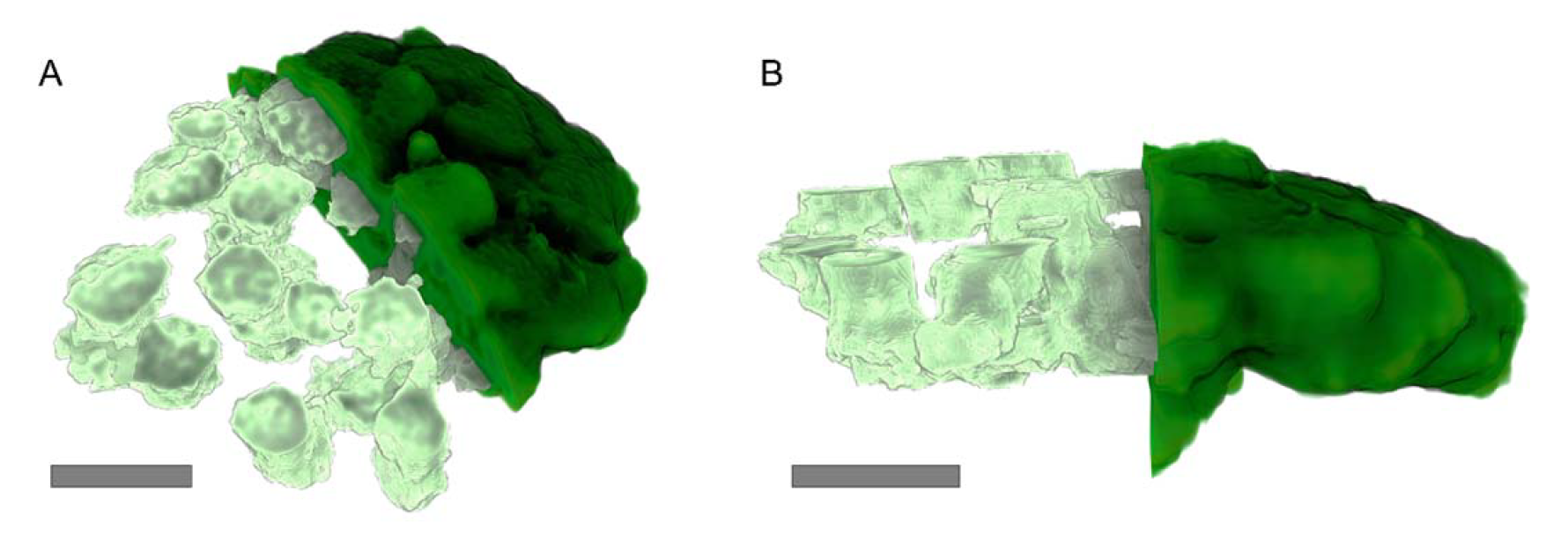
3D reconstruction of a chloroplast (dark green) and its grana (pale green). Grana stacks were segmented by hand from the surroundings. Only half of the complete signal (dark green) and half of the grana is shown. Scalebars represent 1 μm, corrected to indicate pre-expansion dimensions. Images for this reconstruction were made with confocal microscopy.

Typically, chloroplasts in the gel were positioned on their flat side and thus were imaged from the top (figure 3A and B and SI movie 1). This orientation enabled us to accurately determine the shape and diameter of the grana. Grana are well described as ovals and therefore, we could use Stardist, a neural network tool specifically designed to detect round shapes in biological samples, to determine the diameter of the grana (figure 3C-E). We found that there were differences in the size of the grana, even within a single chloroplast (figure 3C and D). The average diameter of the grana was 325 ± 56 nm, but we detected a distribution of grana diameters ranging from 200 to 500 nm (figure 3E, table 2). Additionally, we determined the number of grana per chloroplast to be 91 ± 32 (table 2).

**Figure 3.**
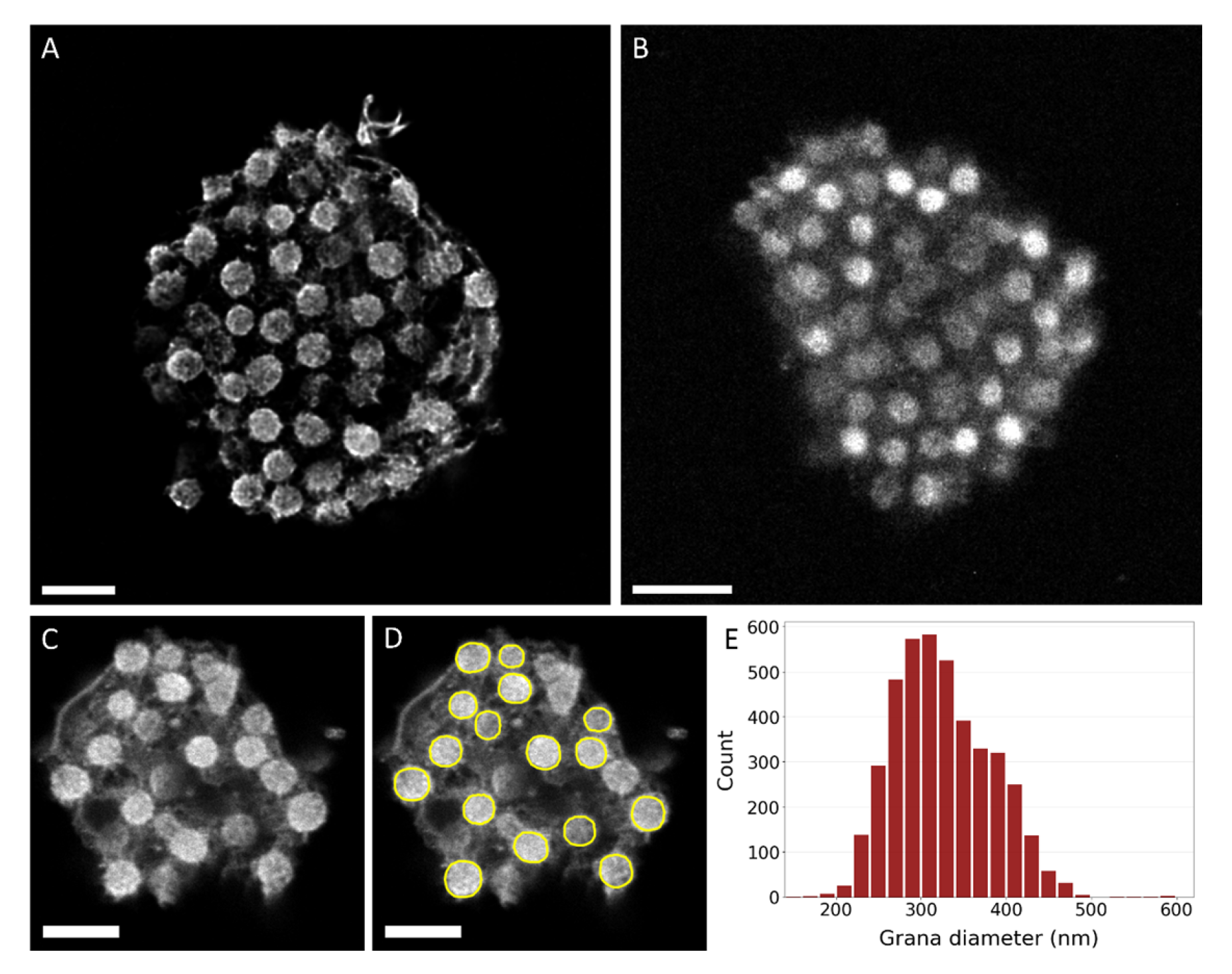
Top-views of expanded chloroplasts. Examples of chloroplasts in top-view as commonly found in the gel, imaged with lattice SIM^2^ (A) or confocal microscopy (B, C). D) Annotation of the grana of the chloroplast from C as performed by Stardist. Scalebars represent 1 μm, corrected to indicate pre-expansion dimensions. E) Histogram of the grana diameter.

**Table 2.**
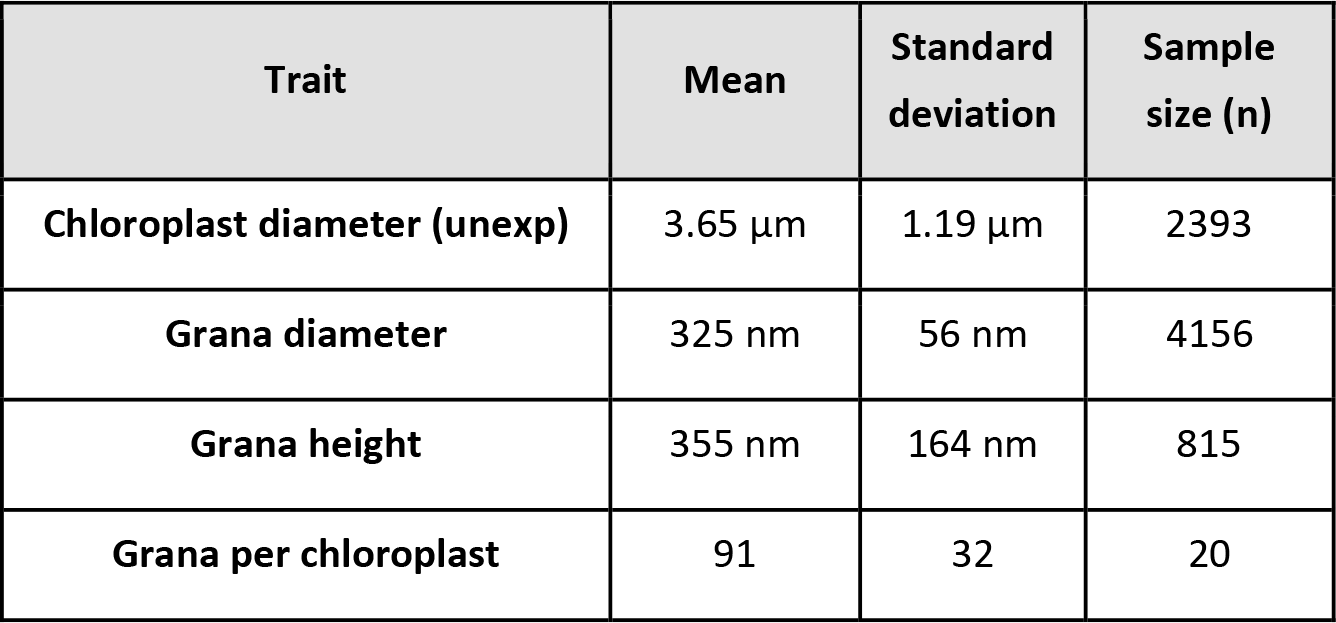
Dimensions of chloroplasts and grana as determined in this study.

In certain instances, we observed chloroplasts lying on their side, providing a side-view (figure 4A and B and SI figure movies 2-3). Using ExM, we imaged chloroplasts that look similar to what is commonly seen with EM, including stroma lamellae appearing on both sides of grana stacks (figure 4C-E). In all occasions, a right-handed wrapping of stroma lamellae around grana stacks was observed (figure 4 and SI figure 3). This is the first observation of the helical wrapping of stroma lamellae using fluorescence microscopy and demonstrates the increased resolution achieved with ExM as compared to fluorescence microscopy. We resolved stroma lamellae that were less than 60 nm apart in pre-expanded dimensions (figure 5). Next, we determined that the height of the grana stacks was 355 ± 164 nm (figure 4F and G). We observed a distribution in stack height, ranging from stacks consisting of only a few membrane layers to grana spanning almost the entire chloroplast.

**Figure 4.**
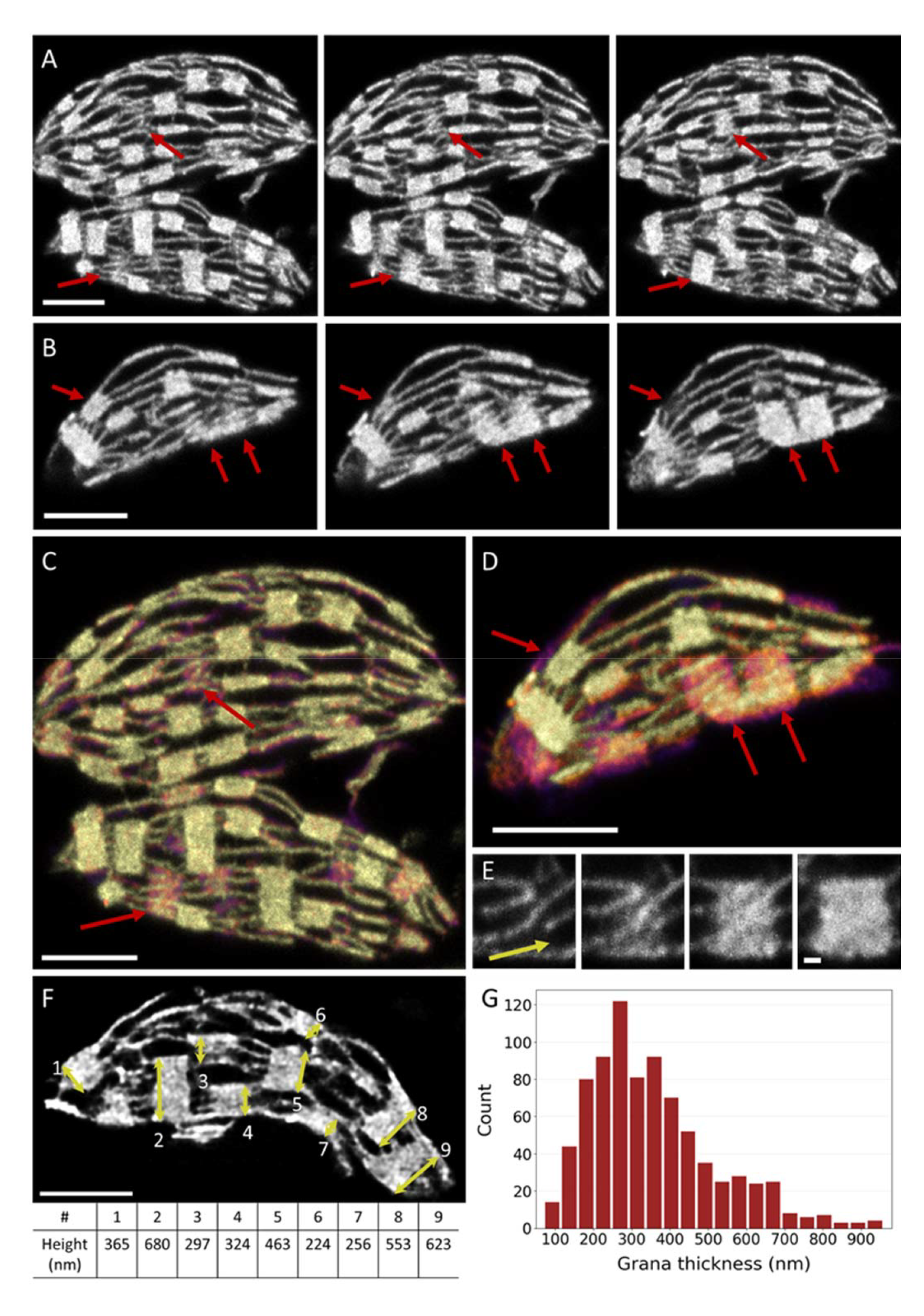
Side-view of expanded chloroplasts. A,B) Montages of side-views of chloroplasts. Red arrows indicate grana where the helical wrapping of the stroma can be observed. Images are 63 nm apart in Z (pre-expansion dimensions). C, D) Depth coded image of the montage in A and B. Red arrows indicate the grana that are also indicated in A and B. E) Example of helical wrapping of the stroma lamellae in a right-handed helix. The yellow arrow shows the wrapping direction of stroma lamellae. F) Example of measurements of grana height. The height of 9 grana stacks was measured and shown in the table. G) Histogram of the grana height. Images were made with confocal microscopy (A-E) or lattice SIM^2^. (F). Scalebar represents 1 μm in A-D and F and 100 nm in E, corrected to indicate pre-expansion dimensions.

**Figure 5.**
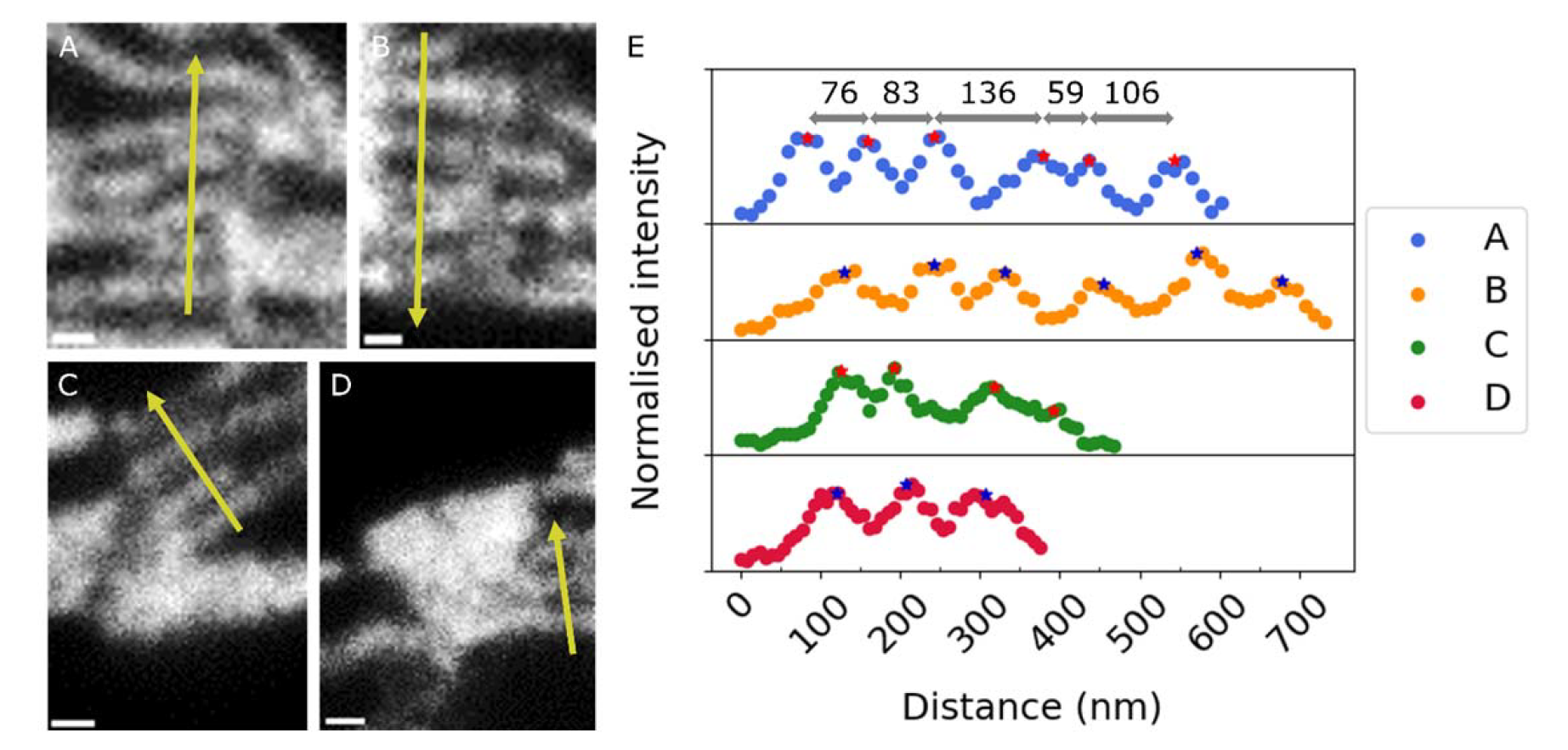
A-D) Examples of stroma lamellae in close proximity to each other, but distinguishable. E) Profile of intensity along the yellow lines in A-D. Peak centers were defined as the middle of the FWHM and are shown with an asterisk. For one profile, the distance between the peak centers is given in nm in pre-expanded dimensions. The minimal distance between the peaks was 59 ± 10 nm in pre-expanded dimensions. Scalebars represent 100 nm in pre-expanded dimensions.

## Discussion

Expansion microscopy - In this work, we used a combination of U-ExM and pan-ExM to image chloroplasts with a more than 5 times improved resolution compared to confocal microscopy. The analysis of top-view and side-view images of chloroplasts resulted in measurements of the grana dimensions with a large sample size (800-4000 measurements) and a high resolution (∼60 nm). Moreover, we presented a 3D model of a complete chloroplast in which grana and stroma lamellae are accurately segmented.

Imaging chloroplasts with ExM offers several advantages over imaging with EM. Firstly, ExM is fast, since it requires only two days of sample preparation and results in a gel full of chloroplasts that can be imaged. Per chloroplast, it takes 1-10 minutes to record its complete structure, thereby outcompeting the speed of EM imaging (Wassie et al., 2019). Moreover, with ExM the volumes that can be imaged are larger. Additionally, the top-view images as obtained with ExM allow for a more direct measurement of the grana width than the cross-section views of EM. Together, that enables us to create large and accurate datasets on the dimensions of the chloroplast and the grana. The resolution reached in this study with ExM is lower than that of EM, but still high enough to accurately determine the diameter of the grana.

A possible drawback of ExM is that isolation, chemical fixation and expansion of chloroplasts might introduce deviations in the thylakoid structure as compared to its native organization. However, on visual inspection, our ExM images look similar to images from EM studies. In particular, the circular shape of grana observed in our top-view images suggests isotropic expansion of the grana. Moreover, the grana diameter we determined with ExM was comparable to literature values for the grana diameter of spinach (325 ± 56 nm vs 325-380 nm for literature, table 2 and 3). However, the height of the grana we determined using ExM differed from literature values (355 ± 164 nm in our measurements vs 91-159 nm for literature, table 2 and 3). This discrepancy can have several causes: 1) The grana height is strongly dependent on the grow light intensity and spectral composition (Anderson et al., 1973; Wagner et al., 2008; Hu et al., 2021). 2) Due to the lower resolution of ExM compared to EM, two grana stacks may have appeared as one in our images and measured accordingly. Furthermore, small grana stacks might not be distinguished from stroma lamellae. 3) The thylakoid membrane could have swollen due to an increased stromal distance and swelling of the lumen, leading to an increased grana height. The thylakoid architecture is dynamic, e.g. light induced swelling of the thylakoid membrane has been reported (Li et al., 2020). Swelling might also occur during isolation and fixation of the de-enveloped chloroplast. In future research projects we aim to develop methods to image the thylakoid organization in intact chloroplast and protoplast. Furthermore, we aim to implement cryo-fixation (Laporte et al., 2022) to assure that the native thylakoid architecture is visualized. Although thylakoid swelling might be a factor, the overall thylakoid macro-organization (number of grana, connection between grana and stroma lamellae, grana diameter) is consistent with EM data, showing that ExM is a suitable technique to study the thylakoid structure and build a 3D model.

Grana dimensions - The grana diameter as determined from the images of expanded chloroplasts (325 ± 56 nm) is comparable to other studies on *S. oleracea* (325-380 nm, table 3). The standard deviation of the grana diameter was 17% of the mean, and a similar distribution is recognized in other studies. In this study, grana ranging from 200-500 nm have been observed. Some outliers might have arisen from inaccuracies of the detection mechanism of the grana by the neural network Stardist, but many of the detected grana have been confirmed by visual inspection. Moreover, we only selected images with a clear grana structure for training and detection of the grana. It has been shown that especially for plants grown in natural light, the distribution in grana diameter is largest (Schumann et al., 2017). In agreement with this, we demonstrated that the grana diameter is far from a fixed value, even in a single chloroplast.

**Table 3.** Dimensions of grana determined with different techniques and on different plant species (Fristedt et al., 2009; Wood et al., 2018; Bussi et al., 2019; Mazur et al., 2019; Wood et al., 2019; Flannery et al., 2021; Sattari Vayghan et al., 2022). The mean value ± standard deviation is given. Where applicable, the values are given for light adapted plants and plants grown in normal light conditions.

| Article | Species | Technique | Grana diameter (nm) | Grana height (nm) | Grana per chloroplast |
| --- | --- | --- | --- | --- | --- |
| Fristedt et al., 2009 | *A. thaliana* | EM | 439 ± 155 | 74 ± 36 | - |
| Wood et al., 2019 | *A. thaliana* | SIM | 350 ± 70 | - | - |
| Mazur et al., 2019 | *A. thaliana* | EM | 600 ± 157 | 156 ± 81 | - |
| Flannery et al., 2021 | *A. thaliana* | SIM | 386 ± 61 | - | - |
| Flannery et al., 2021 | *A. thaliana* | EM | - | 114 ± 69 | - |
| Sattari Vayghan et al., 2022 | *A. thaliana* | EM | 501 ± 143 | 91 ± 39 | - |
| Bussi et al., 2019 | *Lactuca sativa* | EM | 278 ± 43 | 159 ± 94 | - |
| Wood et al., 2018 | *S. oleracea* | EM | 325 ± 74 | 106 ± 64 | - |
| Wood et al., 2018 | *S. oleracea* | SIM | 343 ± 58 | - | 66 ± 9 |
| Wood et al., 2019 | *S. oleracea* | SIM | 380 ± 60 | - | - |

Next to the variation in the grana diameter within a single chloroplast, variation is noticed in the grana diameter reported in literature for 3 different species: *S. oleracea, A. thaliana* and *Lactuca sativa* (table 3). This contrasts studies that have suggested the grana diameter to be conserved between plant species (Albertsson and Andreasson, 2004; Bussi et al., 2019). Many studies have investigated the thylakoid organization of *A. thaliana* and all of these studies report a larger grana diameter than we find for *S. oleracea* (table 3). In addition, the grana diameter reported for *Lactuca sativa* is significantly smaller than ours (unpaired t-test, p<0.0001) (Bussi et al., 2019). A smaller grana diameter is suggested to increase the rate of state transitions, photosystem II (PSII) repair cycle and photosynthetic electron transfer (Wood et al., 2018; Wood et al., 2019; Höhner et al., 2020). Potentially, plants that are more resistant to higher light intensities can benefit from having smaller grana. It should be noted, however, that the light conditions, growth conditions and measuring technique were not the same in the compared literature. All three factors could have influenced the measured grana diameter (Schumann et al., 2017). In fact, Schumann et al. showed that by growing *A. thaliana* in different light intensities, the grana diameter can range from 390 nm (high light) to 570 nm (low light) (Schumann et al., 2017). This range is almost as large as the range of different grana diameters reported in other studies. Therefore, to accurately determine if the grana diameter of plants is significantly different, a single study should compare the grana diameter of different plants species grown in the same light conditions. The same technique to image and detect the grana should be used for all samples. ExM is a suitable technique to investigate this kind of differences in thylakoid build-up between species.

Improvements - Although ExM is a fast and accurate imaging technique, we believe that ExM on chloroplasts can be improved to become more reliable and versatile. Most importantly, fixation of the chloroplasts was not successful in all attempts, which led to expanded chloroplasts without clear grana structure. We often observed expanded chloroplasts without grana structure when we used plants grown in a phytotron. The protocol that was used in this study was optimized for spinach purchased from the local grocer. Potentially, the isolation and fixation of chloroplasts from outdoor grown plants was more successful, because the thylakoid organization of plants grown in natural sunlight is different to those grown in artificial light (Schumann et al., 2017). Furthermore, the variable conditions outdoor could make these plants sturdier and more suitable for the preparation for ExM. The protocol for fixation should be optimized to make it better applicable on other plant species and plants grown in a phytotron. Furthermore, a method should be developed that assures that the fully native thylakoid architecture is visualized. Taken together this will allow to study light adaptation responses of the thylakoid membrane and to use mutant variants of the model plant *A. thaliana*.

Future perspective – The application of ExM in the field of photosynthesis offers a promising avenue to address several outstanding questions. ExM can facilitate the intuitive visualization of the folding of thylakoid membranes through the construction of a 3D model of the entire chloroplast. In addition, the dimensions of grana can be readily determined from a large dataset obtained with ExM, thereby aiding in the investigation of differences in thylakoid structure. Specifically, ExM can be employed to study the thylakoid architecture of different species or adaptations to varying light conditions. Next, ExM can be combined with antibody staining and hence enable accurate and fast protein localization (Gambarotto et al., 2019). Typically, antibody staining is challenging in chloroplasts due to limited antibody diffusion into the appressed regions and high background fluorescence from chlorophyll. With ExM, the sample is de-crowded and chlorophylls are washed away, which lowers the background and creates space in the appressed regions for primary and secondary antibodies. This technique could allow the staining and localization of key photosynthetic proteins, such as PSII and light harvesting complex II (LHCII), and facilitate the tracking of their location during different stages of state transitions or after high light damage. Furthermore, ExM could potentially enable the use of single molecule or super resolution imaging techniques on the thylakoid membrane by reducing background fluorescence from chlorophyll (Gambarotto et al., 2019). Combining high-resolution data on the thylakoid structure with the location of key photosynthetic proteins could provide valuable insights into the link between protein composition and thylakoid ultrastructure.

## Acknowledgements

This work was supported by the Dutch Organisation for Scientific Research (NWO) via a Vidi grant no. VI.Vidi 192.042 (E.W.). The authors would like to thank Francesco Saccon for critically reading the manuscript. Additionally, we would like to express our appreciation to Dr. Edwin Lamers (Carl Zeiss bv) and Dr. Abel Pereira da Graça (Carl Zeiss Microscopy GmbH) for the opportunity to use the ZEISS Elyra 7 with Lattice SIM² and their technical assistance.

## References

Albertsson PÅ, Andreasson E (2004) The constant proportion of grana and stroma lamellae in plant chloroplasts. Physiologia Plantarum 121: 334–342

Anderson JM, Goodchild D, Boardman N (1973) Composition of the photosystems and chloroplast structure in extreme shade plants. Biochimica et Biophysica Acta (BBA)-Bioenergetics 325: 573–585

Anderson JM, Horton P, Kim E-H, Chow WS (2012) Towards elucidation of dynamic structural changes of plant thylakoid architecture. Philosophical Transactions of the Royal Society B: Biological Sciences 367: 3515–3524

Armbruster U, Labs M, Pribil M, Viola S, Xu W, Scharfenberg M, Hertle AP, Rojahn U, Jensen PE, Rappaport F (2013) Arabidopsis CURVATURE THYLAKOID1 proteins modify thylakoid architecture by inducing membrane curvature. The Plant Cell 25: 2661–2678

Blankenship RE (2021) Molecular mechanisms of photosynthesis. John Wiley & Sons

Bussi Y, Shimoni E, Weiner A, Kapon R, Charuvi D, Nevo R, Efrati E, Reich Z (2019) Fundamental helical geometry consolidates the plant photosynthetic membrane. Proceedings of the National Academy of Sciences 116: 22366–22375

Caffarri S, Kouřil R, Kereïche S, Boekema EJ, Croce R (2009) Functional architecture of higher plant photosystem II supercomplexes. The EMBO journal 28: 3052–3063

Chen F, Tillberg PW, Boyden ES (2015) Expansion microscopy. Science 347: 543–548

Damstra HG, Mohar B, Eddison M, Akhmanova A, Kapitein LC, Tillberg PW (2022) Visualizing cellular and tissue ultrastructure using Ten-fold Robust Expansion Microscopy (TREx). Elife 11: e73775

Engel BD, Schaffer M, Kuhn Cuellar L, Villa E, Plitzko JM, Baumeister W (2015) Native architecture of the Chlamydomonas chloroplast revealed by in situ cryo-electron tomography. elife 4: e04889

Flannery SE, Hepworth C, Wood WH, Pastorelli F, Hunter CN, Dickman MJ, Jackson PJ, Johnson MP (2021) Developmental acclimation of the thylakoid proteome to light intensity in Arabidopsis. The Plant Journal 105: 223–244

Fristedt R, Willig A, Granath P, Crevecoeur M, Rochaix J-D, Vener AV (2009) Phosphorylation of photosystem II controls functional macroscopic folding of photosynthetic membranes in Arabidopsis. The Plant Cell 21: 3950–3964

Gambarotto D, Zwettler FU, Le Guennec M, Schmidt-Cernohorska M, Fortun D, Borgers S, Heine J, Schloetel J-G, Reuss M, Unser M (2019) Imaging cellular ultrastructures using expansion microscopy (U-ExM). Nature methods 16: 71–74

Gómez-de-Mariscal E, García-López-de-Haro C, Ouyang W, Donati L, Lundberg E, Unser M, Muñoz-Barrutia A, Sage D (2021) DeepImageJ: A user-friendly environment to run deep learning models in ImageJ. Nature Methods 18: 1192–1195

Höhner R, Pribil M, Herbstová M, Lopez LS, Kunz H-H, Li M, Wood M, Svoboda V, Puthiyaveetil S, Leister D (2020) Plastocyanin is the long-range electron carrier between photosystem II and photosystem I in plants. Proceedings of the National Academy of Sciences 117: 15354–15362

Hu C, Nawrocki WJ, Croce R (2021) Long-term adaptation of Arabidopsis thaliana to far-red light. Plant, cell & environment 44: 3002–3014

Hu Y, Limaye A, Lu J (2020) Three-dimensional segmentation of computed tomography data using Drishti Paint: new tools and developments. Royal Society Open Science 7: 201033

Iwai M, Roth MS, Niyogi KK (2018) Subdiffraction-resolution live-cell imaging for visualizing thylakoid membranes. The Plant Journal 96: 233–243

Johnson MP, Vasilev C, Olsen JD, Hunter CN (2014) Nanodomains of cytochrome b 6 f and photosystem II complexes in spinach grana thylakoid membranes. The Plant Cell 26: 3051–3061

Johnson MP, Wientjes E (2020) The relevance of dynamic thylakoid organisation to photosynthetic regulation. Biochimica et Biophysica Acta (BBA)-Bioenergetics 1861: 148039

Kaftan D, Brumfeld V, Nevo R, Scherz A, Reich Z (2002) From chloroplasts to photosystems: in situ scanning force microscopy on intact thylakoid membranes. The EMBO journal 21: 6146–6153

Kirchhoff H (2014) Diffusion of molecules and macromolecules in thylakoid membranes. Biochimica et Biophysica Acta (BBA)-Bioenergetics 1837: 495–502

Lai HM, Liu AKL, Ng W-L, DeFelice J, Lee WS, Li H, Li W, Ng HM, Chang RC-C, Lin B (2016) Rationalisation and validation of an acrylamide-free procedure in three-dimensional histological imaging. PLoS One 11: e0158628

Laporte MH, Klena N, Hamel V, Guichard P (2022) Visualizing the native cellular organization by coupling cryofixation with expansion microscopy (Cryo-ExM). Nature methods 19: 216–222

Li M, Mukhopadhyay R, Svoboda V, Oung HMO, Mullendore DL, Kirchhoff H (2020) Measuring the dynamic response of the thylakoid architecture in plant leaves by electron microscopy. Plant Direct 4: e00280

Limaye A (2012) Drishti: a volume exploration and presentation tool. *In* Developments in X-ray Tomography VIII, Vol 8506. SPIE, pp 191–199

Liu L-N, Scheuring S (2013) Investigation of photosynthetic membrane structure using atomic force microscopy. Trends in plant science 18: 277–286

M’Saad O, Bewersdorf J (2020) Light microscopy of proteins in their ultrastructural context. Nature communications 11: 1–15

Mazur R, Mostowska A, Szach J, Gieczewska K, Wójtowicz J, Bednarska K, Garstka M, Kowalewska Ł (2019) Galactolipid deficiency disturbs spatial arrangement of the thylakoid network in Arabidopsis thaliana plants. Journal of Experimental Botany

Mehta M, Sarafis V, Critchley C (1999) Thylakoid membrane architecture. Functional Plant Biology 26: 709–716

Mustardy L, Buttle K, Steinbach G, Garab Gz (2008) The three-dimensional network of the thylakoid membranes in plants: quasihelical model of the granum-stroma assembly. The Plant Cell 20: 2552–2557

Mustárdy L, Garab G (2003) Granum revisited. A three-dimensional model–where things fall into place. Trends in plant science 8: 117–122

Onoa B, Fukuda S, Iwai M, Bustamante C, Niyogi KK (2020) Atomic force microscopy visualizes mobility of photosynthetic proteins in grana thylakoid membranes. Biophysical journal 118: 1876–1886

Paolillo Jr DJ (1970) The three-dimensional arrangement of intergranal lamellae in chloroplasts. Journal of Cell Science 6: 243–253

Pribil M, Labs M, Leister D (2014) Structure and dynamics of thylakoids in land plants. Journal of experimental botany 65: 1955–1972

Sattari Vayghan H, Nawrocki WJ, Schiphorst C, Tolleter D, Hu C, Douet V, Glauser G, Finazzi G, Croce R, Wientjes E (2022) Photosynthetic light harvesting and thylakoid organization in a CRISPR/Cas9 Arabidopsis thaliana LHCB1 knockout mutant. Frontiers in plant science 13: 833032

Schmidt U, Weigert M, Broaddus C, Myers G (2018) Cell detection with star-convex polygons. *In* International Conference on Medical Image Computing and Computer-Assisted Intervention. Springer, pp 265-273

Schumann T, Paul S, Melzer M, Dörmann P, Jahns P (2017) Plant growth under natural light conditions provides highly flexible short-term acclimation properties toward high light stress. Frontiers in plant science 8: 681

Shimoni E, Rav-Hon O, Ohad I, Brumfeld V, Reich Z (2005) Three-dimensional organization of higher-plant chloroplast thylakoid membranes revealed by electron tomography. The Plant Cell 17: 2580–2586

Wagner R, Dietzel L, Bräutigam K, Fischer W, Pfannschmidt T (2008) The long-term response to fluctuating light quality is an important and distinct light acclimation mechanism that supports survival of Arabidopsis thaliana under low light conditions. Planta 228: 573–587

Wassie AT, Zhao Y, Boyden ES (2019) Expansion microscopy: principles and uses in biological research. Nature methods 16: 33–41

Weigert M, Schmidt U, Haase R, Sugawara K, Myers G (2020) Star-convex polyhedra for 3D object detection and segmentation in microscopy. In Proceedings of the IEEE/CVF Winter Conference on Applications of Computer Vision, pp 3666–3673

Wietrzynski W, Schaffer M, Tegunov D, Albert S, Kanazawa A, Plitzko JM, Baumeister W, Engel BD (2020) Charting the native architecture of Chlamydomonas thylakoid membranes with single-molecule precision. Elife 9: e53740

Wildman S, Hirsch AM, Kirchanski S, Spencer D (2005) Chloroplasts in living cells and the string-of-grana concept of chloroplast structure revisited. Discoveries in Photosynthesis: 737–744

Wood WH, Barnett SF, Flannery S, Hunter CN, Johnson MP (2019) Dynamic thylakoid stacking is regulated by LHCII phosphorylation but not its interaction with PSI. Plant physiology 180: 2152–2166

Wood WH, MacGregor-Chatwin C, Barnett SF, Mayneord GE, Huang X, Hobbs JK, Hunter CN, Johnson MP (2018) Dynamic thylakoid stacking regulates the balance between linear and cyclic photosynthetic electron transfer. Nature plants 4: 116–127

Zhang C, Kang JS, Asano SM, Gao R, Boyden ES (2020) Expansion microscopy for beginners: visualizing microtubules in expanded cultured HeLa cells. Current protocols in neuroscience 92: e96

